# A genetic driver of epileptic encephalopathy impairs gating of synaptic glycolysis

**DOI:** 10.1101/2025.06.17.660213

**Authors:** Zhanat Koshenov, Alexandros C. Kokotos, Lorena Benedetti, Jennifer Lippincott Schwartz, Timothy A. Ryan

## Abstract

The brain is a disproportionately large consumer of fuel, estimated to expend ∼20% of the whole-body energy budget, and therefore it is critical to adequately control brain fuel expenditures while satisfying its on-demand needs for continued function. The brain is also metabolically vulnerable as the inability to adequately fuel cellular processes that support information transfer between cells leads to rapid neurological impairment. We show here that a genetic driver of early onset epileptic encephalopathy (EOEE), SLC13A5, a Na^+^/citrate cotransporter (NaCT), is critical for gating the activation of local presynaptic glycolysis. We show that SLC13A5 is in part localized to a presynaptic pool of membrane-bound organelles and acts to transiently clear axonal citrate during electrical activity, in turn activating phosphofructokinase 1. We show that loss of SLC13A5 or mistargeting to the plasma membrane results in suppressed glycolytic gating, activity dependent presynaptic bioenergetic deficits and synapse dysfunction.

**Graphical abstract:** 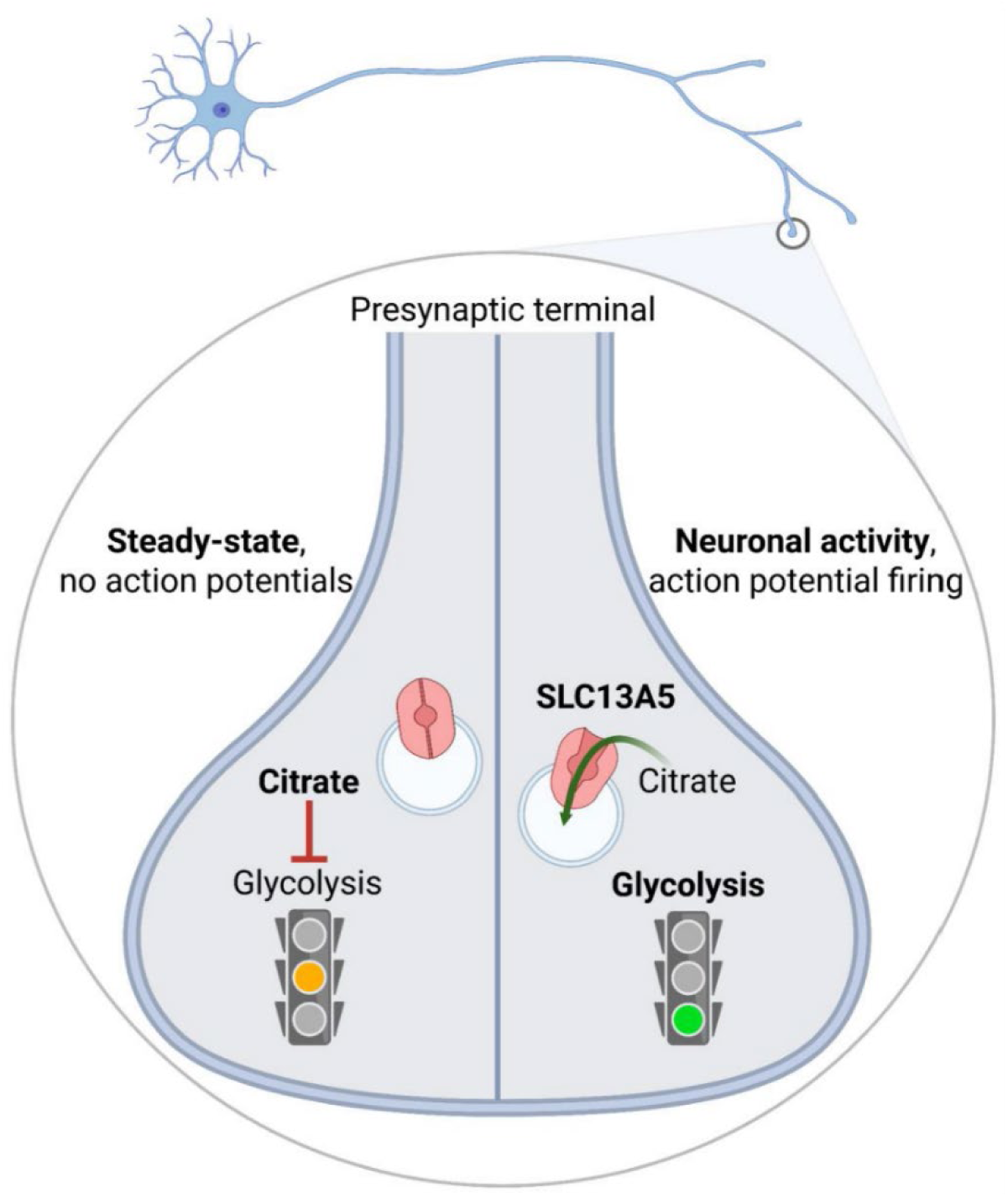

## Introduction

Our brains are metabolically vulnerable organs and maintaining the proper balance between fuel use and availability is critical for brain function. In humans, a 50% decrease in plasma glucose leads to the rapid onset of neurological symptoms including delirium, seizures and coma^1^. This vulnerability reveals that the need for a combustible carbon source for many neural circuits is easily outstripped when fuel delivery is curtailed and rapid conversion of an appropriate carbon source into the key biochemical currency, ATP, is required to sustain brain function. This acute vulnerability has deep relevance to neurological disease. Brain hypometabolism is considered one of the first signs of the onset of dementias both in sporadic Alzheimer’s disease (AD)^2,3^ and Parkinson’s disease (PD)^4–6^ and drivers of brain hypoglycemia, such as Glut-1 deficiency, are tightly linked to EOEE^7^. Genetic ablation of SLC13A5 in mice^8^ and fish^9^ leads to seizure activity, but loss of function in flies, where it is primarily expressed in oenocytes and the gut, prolongs lifespan^10^. The basis for SLC13A5 deficiency driving EOEE in humans has remained enigmatic, owing to the lack of understanding of this protein’s function in neuronal tissue. In general, SCL13A5 is considered to be a plasma membrane transporter, as its inhibition in hepatocytes and heterologous cells reduces extracellular citrate uptake^11–13^, while loss of function in liver protects against high fat diet driven dyslipidemia^14^ likely associated with a change in the control of hepatic fatty acid synthesis. We and others have previously shown that nerve terminals are enriched in glycolytic machinery^15,16^ that this machinery is used to rapidly upregulate local ATP production during bursts of electrical activity^17^, but failure to do so leads to rapid collapse in synaptic function^18^. Although substrate and product feedback inhibition influence several steps in glycolysis, phosphofructokinase 1 (PFK1) is known across different tissues to be controlled via both second messengers and metabolites, including citrate^19^. In muscle, robust activity is known to downregulate PFK1 function via the accumulation of citrate derived from the mitochondrial tricarboxylic cycle (TCA) ^20,21^.We show here that electrical activity gates presynaptic glycolysis by triggering the activation of PFK1. A transient burst of action potential (AP) firing leads to a synchronous burst of fructose-1,6-bisphosphate (FBP), PFK1’s product. We show here that AP driven activation of PFK1 at nerve terminals is due to a transient clearance of cytoplasmic citrate, that is mediated by SLC13A5. As SLC13A5 is a Na^+^ dependent citrate co-transporter, copies of this on the plasma membrane would not be capable of exporting citrate since the Na^+^ ion gradient would always favor influx. We demonstrate here that, in contrast with hepatocytes, neurons show minimal citrate uptake when perfused with citrate but that an intracellular pool of SCL13A5 is responsible for citrate clearance during electrical activity and the subsequent activation of PFK1. Our experiments demonstrate that chronic loss of SLC13A5 in axons leads to diminished presynaptic ATP that can be restored by silencing electrical activity. We show that the loss of proper PFK1 activation in turn results in the failure to adequately synthesize sufficient ATP, resulting in activity-dependent ATP deficits, slowed recovery of presynaptic ATP levels following bursts of activity and degradation of synaptic vesicle recycling kinetics. We further find that the bioenergetic deficits at nerve terminals resulting from loss of SLC13A5 can be reversed by approaches that activate glycolysis, including the use of Terazosin, an FDA-approved treatment for benign prostate hyperplasia that has also been shown to activate phosphoglycerate kinase-1, the first ATP producing enzyme in glycosis^16,22,23^. These results help offer a mechanistic basis for how loss of SLC13A5 leads to neurological impairment and offers a promising therapeutic direction to alleviate suffering from loss of SLC13A5.

## Results

### SLC13A5 is important for activity dependent presynaptic ATP production

Given the neurological manifestation of SLC13A5 mutations^24^ and the known control of glycolysis by citrate^25^, we asked whether this NaCT is important for synaptic bioenergetics using a recently developed genetically encoded ATP biosensor targeted to nerve terminals^26^ (Figure 1 a). Transient knock down (KD) of SLC13A5 in rat hippocampal neurons (Figure S1 a-d) resulted in nearly 60% reduction of basal presynaptic ATP (Figure 1 b). This bioenergetic deficit could be rescued by wild type (WT) human transporter, but not the transporter with a common disease-causing glycine 219 to arginine mutation ^27^ (Figure 1 b). To test if the ATP deficit caused by downregulation of SLC13A5 is dependent on neuronal activity, we used tetrodotoxin (TTX), a Na^+^ channel blocker, to inhibit spontaneous neuronal activity in culture for several days before the measurement. TTX treatment equilibrated presynaptic ATP of SLC13A5 KD neurons to that of control neurons (Figure 1 c), suggesting activity dependence of SLC13A5’s role in synaptic bioenergetics. To directly test this claim, we examined the dynamics of synaptic ATP during stimulation with an extended train of action potentials (AP) at 10 Hz after removal of TTX. These experiments demonstrate although such bursts of activity lead to transient drop of presynaptic ATP in both control and SLC13A5 KD nerve terminals, loss of SLC13A5 leads to much slower recovery of ATP levels post-stimulation (Figure 1 d, e). Similar results were obtained in cells that were not pretreated with TTX: controls were able to recover presynaptic ATP post-stimulation, while SLC13A5 KD neurons had larger percentage ATP drop during stimulation that failed to recover ATP in the post-stimulus period. These deficits were rescued by expression of an shRNA-insensitive WT variant of SLC13A5, but not with the G129R mutant (Figure S1 e-g). We used a single metric to characterize the net changes in ATP levels during and after electrical activity. On average control neurons show a net increase in ATP levels 150 sec after prolonged electrical stimulation while loss of SLC13A5 results in a net loss of ATP (Figure S1 f,g).

**Figure 1.**
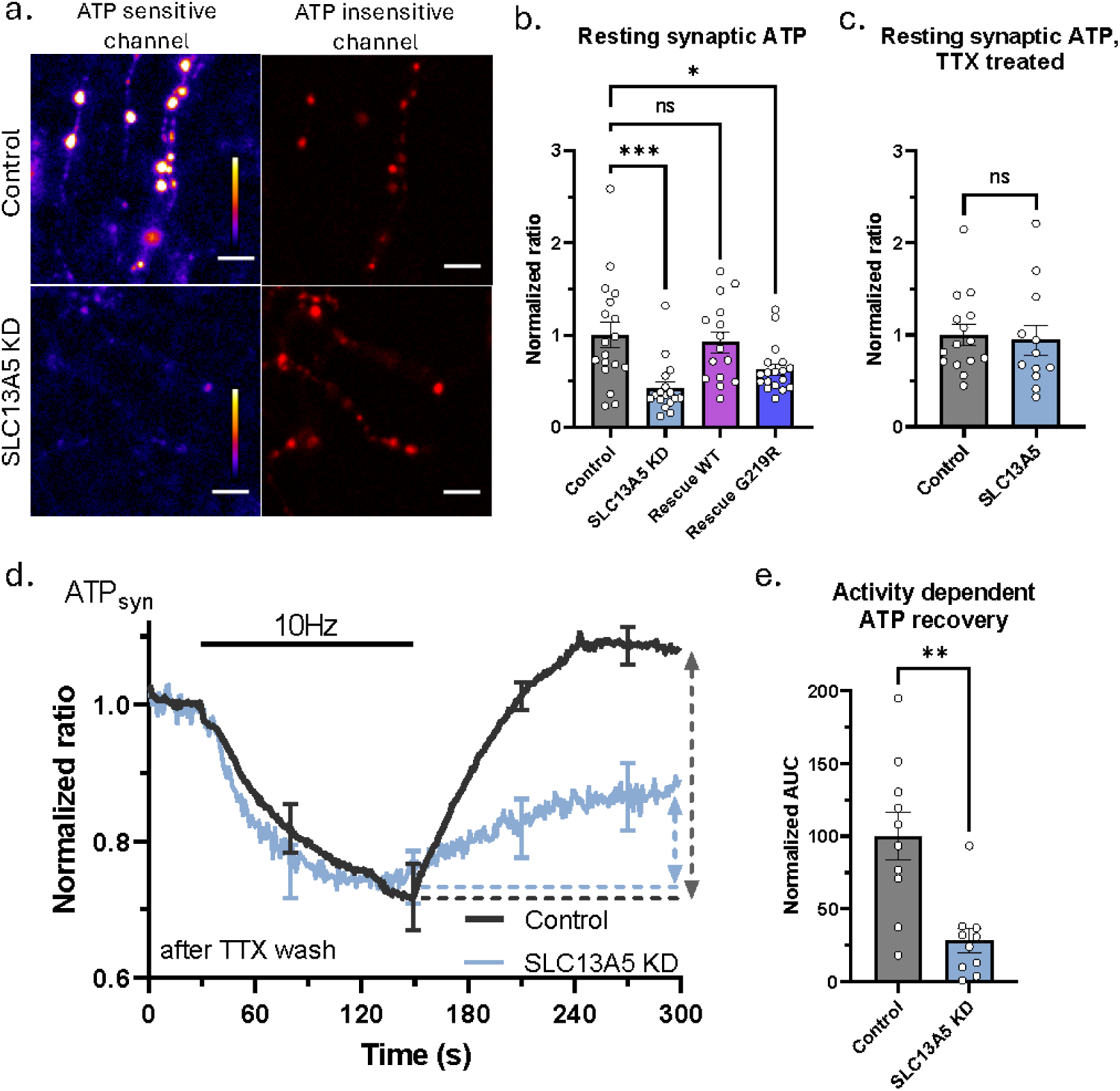
SLC13A5 is important for activity dependent presynaptic ATP production. (a) Representative images of control and SLC13A5 KD hippocampal neurons expressing synapto-iATPSnFR2-miRFP670nano3. ATP sensitive channel is pseudo-colored with calibration bar ranging from 0 to 100 AFU. Scale bar is 5 µm. (b) Mean resting presynaptic ATP +/-SEM depicted as iATPSnFR2-miRFP670nano3 ratio, normalized to control. Control, n=18; SLC13A5 KD, n=17; Rescue with WT SLC13A5, n=15; Rescue with G219R mutant of SLC13A5, n=18. (c) Mean resting presynaptic ATP +/-SEM of TTX treated neurons, normalized to control ratio. Control, n=15; SLC13A5 KD, n=12. (d) Average +/-SEM synapto-iATPSnFR2-miRFP670nano3 traces for control (black) and SLC13A5 KD (light blue) neurons stimulated with 1200 APs at 10 Hz. Dashed lines and arrows show area quantified in e. (e) Mean activity dependent presynaptic ATP recovery +/-SEM, quantified as AUC after the end of AP train (shown with dashed lines and arrows in d), n=10 for both.

In line with activity dependence of SLC13A5 function, acute inhibition of SLC13A5 with a known inhibitor, PF-06649298 ^28^, did not impact steady-state presynaptic ATP (Figure S1 h). Acute electrical stimulation, on the other hand, resulted in diminished net activity dependent ATP change after inhibitor treatment (Figure S1 i, j). To gain insight into preferential involvement of SLC13A5 in mitochondrial versus glycolytic ATP production pathways, we substituted glucose with lactate and pyruvate. When using mitochondrial fuels, neurons had identical net activity dependent ATP change before and after inhibitor treatment with or without SLC13A5 inhibition (Figure S1 k, l). These experiments support the notion that SLC13A5 plays an important role in activity dependent ATP production at nerve terminals and suggest that its mechanism of action likely involves glycolysis.

### Neuronal activity drives presynaptic glycolysis that depends on SLC13A5 and citrate sensitivity of PFK1

To understand the role of the NaCT in presynaptic glycolysis we made use of recently developed genetically encoded fructose-1,6-bisphosphate (FBP) biosensor^29^ (Figure 2) targeted to nerve terminals to track the activity of presynaptic PFK1, a key glycolytic enzyme that is known to be regulated by citrate^21,25^. These experiments showed that a brief burst of action potential firing led to a transient increase in presynaptic FBP (Figure 2 b, c) that decayed back to resting levels over the next minute. These data suggest electrical activity activates PFK1 at presynaptic terminals consistent with a previous report of activity dependent boost in neuronal glycolysis^30^. Given the known role of citrate as an allosteric inhibitor of PFK1^19^, and the impact of loss of SLC13A5 on activity driven bioenergetics (Figure 1), we examined the impact of shRNA-mediated KD of SLC13A5 of the observed activity-driven PFK1 activation. These experiments show that loss of SLC13A5 reduced the maximum activity dependent FBP peak by ∼ 50% (Figure 2 b, c), without altering the resting levels of FBP (Figure S2 a), revealing that the transporter plays a role in presynaptic activity dependent glycolysis.

**Figure 2.**
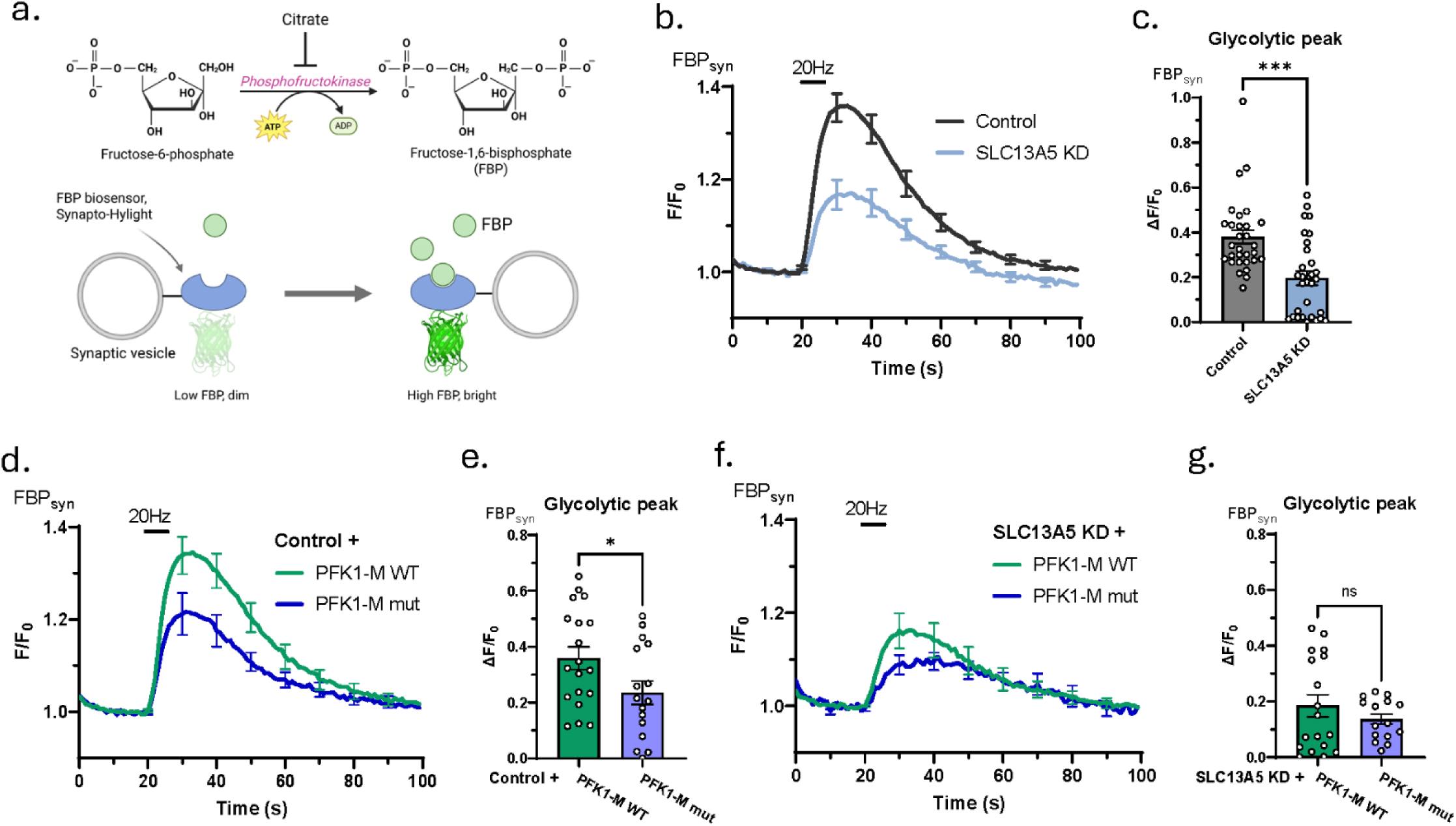
Neuronal activity drives presynaptic glycolysis that depends on SLC13A5 and citrate sensitivity of PFK1. (a) Schematic illustrations of enzymatic reaction catalyzed by PFK1 (upper panel) and mechanism of action of FBP biosensor, synapto-Hylight’s (lower panel). (b, d, f) Average +/-SEM synapto-Hylight traces for (b) control (black) and SLC13A5 KD (light blue), (d) control PFK1-M WT OE (green) and control PFK1-M mut OE (dark blue), (f) SLC13A5 KD PFK1-M WT OE (green) and SLC13A5 PFK1-M mut OE (dark blue) neurons stimulated with 100 APs at 20 Hz. (c, e, g) Mean glycolytic peak (maximal synapto-Hylight signal increase over baseline) +/-SEM in response to 100 APs at 20 Hz, quantified for (c) control (n=30) and SLC13A5 KD (n=30), (e) control PFK1-M WT OE (n=19) and control PFK1-M mut OE (n=16), (g) SLC13A5 KD PFK1-M WT OE (n=18) and SLC13A5 PFK1-M mut OE (n=12) neurons.

Citrate is known to directly inhibit PFK1 through an allosteric binding site and mutations in this binding site dramatically alter the negative allosteric regulation of this enzyme ^25^. In order to determine if citrate plays a role in PFK1 activation at nerve terminals, we examined presynaptic FBP dynamics triggered by electrical activity in hippocampal nerve terminals in which we either overexpressed a mutant isoform of PFK1 (PFK1-M mut) with reduced citrate sensitivity^25^ or a WT PFK1-M as a control. Expression of the PFK1-M mut with reduced citrate sensitivity resulted in diminished glycolytic activation compared to WT PFK1-M overexpressing neurons (Figure 2 d, e), without altering resting FBP levels (Figure S2 b), supporting the idea that activity-driven activation of PFK1 is modulated in part by changes in cytosolic citrate levels. In contrast, in SLC13A5 KD neurons, expression of either WT or mutant PFK1 did not impact glycolytic activation by electrical activity or resting FBP levels (Figure 2 f, g; Figure S2 b, c). Overall, these results show that neuronal activity triggers glycolytic activation through citrate dependent regulation of PFK1 that relies on SLC13A5.

### Neuronal SLC13A5 functions on an internal pool of slowly recycling vesicles

Humans suffering from SLC13A5 deficiency have elevated plasma citrate levels supporting the idea that loss of this NaCT function results from diminished citrate uptake across the plasma membrane ^31^. In hepatocytes, acute pharmacological block of SLC13A5 leads to impaired citrate uptake, supporting the idea that liver NaCT acts on the plasma membrane. In order to determine if SLC13A5’s impact on axonal bioenergetics was due to a potential problem with plasma membrane citrate uptake in axons, we made use of genetically encoded citrate sensor, Citroff1 ^32^, to determine if acute application of extracellular citrate would lead to increases in cytoplasmic citrate in the axons of primary neurons, and if so, if it depends on SLC13A5. Acute application of 1 mM extracellular citrate led to only small changes in cytoplasmic citrate (Fig. 3 a, b) that was not statistically different than that measured in SLC13A5 KD axons. Although this signal was quite small and could in principle be driven by changes such as pH alterations, use of the citrate-insensitive sensor that has similar pH-dependence^32^ (Fig. 3 a, b) showed that even this small signal is not likely due to a pH artifact. In contrast to these small signals driven by citrate superfusion in WT neurons, overexpression of SLC13A5 harboring an N-terminal fluorescent protein tag led to ∼ 23-fold increase in axonal citrate compared to WT neurons, while expression of a c-terminal tagged SLC13A5 led to only a ∼3-fold increase and that of an untagged SLC13A5 led to a ∼7-fold increase (Figure 3 c, d; Figure S3 a). Given the wide dynamic range (3-23 fold compared to WT) in extracellular citrate uptake observed with these different constructs, we asked if there was any difference in their ability to rescue the SLC13A5 KD mediated suppression of glycolytic activation during action potential firing. Remarkably these experiments demonstrate that despite the ability to mediate very strong plasma membrane citrate uptake, the N-terminal construct failed to restore glycolytic activation while the C-terminal tagged variant fully rescued the loss of SLC13A5 (Figure 3 e, f). These data strongly suggest that in WT neurons there is relatively little expression of SLC13A5 on the plasma membrane, and that instead it functions from an intracellular pool.

**Figure 3.**
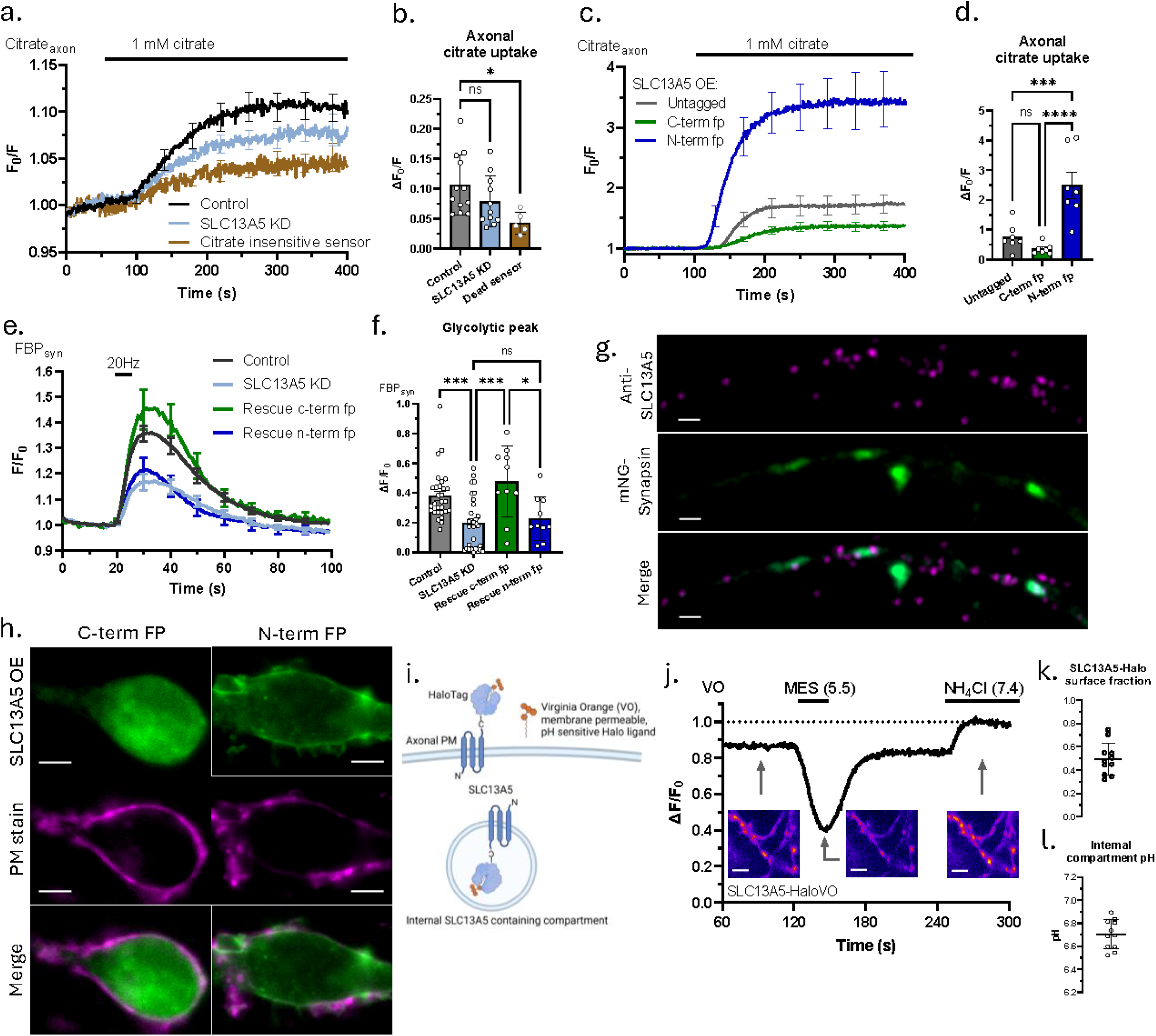
Neuronal SLC13A5 functions in an internal pool of slowly recycling vesicles. (a, c) Average +/-SEM axonal Citroff1 traces in response to 1 mM citrate superfusion in (a) control (black, n=12), SLC13A5 KD (light blue, n=12) and citrate insensitive sensor (CitroffRH) expressing neurons (brown, n=5), and (c) neurons overexpressing untagged (grey, n=7), c-terminally tagged (green, n=7), and n-terminally tagged (dark blue, n=7) SLC13A5 fusion constructs. (b, d) Mean axonal citrate uptake (maximal Citroff1 signal increase over baseline) +/-SEM in response to 1 mM citrate superfusion, quantified for (b) control (n=12), SLC13A5 KD (n=12) and CitroffRH expressing neurons (n=5), and (d) neurons overexpressing untagged (n=7), c-terminally tagged (n=7), and n-terminally tagged (n=7) SLC13A5 fusion constructs. (e) Average +/-SEM synapto-Hylight traces for control (black), SLC13A5 KD (light blue), rescue with c-terminally tagged hSLC13A5 (green), and rescue with n-terminally tagged hSLC13A5 (dark blue) neurons stimulated with 100 APs at 20 Hz. (f) Mean glycolytic peak +/-SEM in response to 100 APs at 20 Hz, quantified for control (n=30), SLC13A5 KD (n=30), rescue with c-terminally tagged hSLC13A5 (n=10), and rescue with n-terminally tagged hSLC13A5 (n=10) neurons. (g) Representative high-resolution Airyscan images of an axonal region showing endogenous SLC13A5 (upper panel), mNeonGreen-Synapsin (middle panel), and merged image (lower panel). Scale bar is 1 µm. (h) Representative images of c-, and n-terminally mEmerald-tagged SLC13A5 fusion proteins co-labelled with CellMask Orange PM stain. Scale bar is 5 µm. (i) Schematic illustration of predicted intracellular localization of SLC13A5-Halo construct and labelling with VO dye. (j) Representative VO trace in response to short acid and later NH_4_Cl superfusion. Inserts are images of axonal regions corresponding to indicated stages of the experimental protocol. Scale bar is 5 µm. (k) Mean surface fraction +/-SEM of SLC13A5-Halo construct, n=11 (l) Mean pH +/-SEM of internal SLC13A5-Halo containing compartment, n=11

To elucidate the intracellular localization of the transporter in neurons, we performed high-resolution imaging of endogenous SLC13A5 using specific antibodies and OE of fluorescently tagged transporters described above. In neuronal processes, endogenous SLC13A5 showed a sparse punctate distribution that partially overlapped with presynaptic compartments (Figure 3 g). In neuronal somas, endogenous SLC13A5 was localized in the trans-Golgi network (TGN), evident from its co-localization with TGN resident protein, sialyltransferase^33^ (Figure S3 b).

Curiously, overexpression of human SLC13A5 fusion protein resulted in differential localization depending on the position of the fluorescent tag. While tagging the transporter from its C-terminal resulted in intracellular distribution similar to that of the endogenous protein, tagging SLC13A5 from its N-terminal resulted in its preferential localization on the plasma membrane (PM) (Figure 3 h). Both fusion proteins were present in axons, although their possible differential distribution was difficult to discern (Figure S3 c).

Since C-terminally tagged SLC13A5 construct functionally mimics the endogenous transporter, we used it to further investigate its axonal localization. Based on available structures^12^, the C-terminus of SLC13A5 will face the extracellular space when it resides on the PM and in the lumen of an intracellular compartment when localized on internal membranes (Figure S3 d). Using HaloTag^34^ as a C-terminal fusion protein for SLC13A5, we applied a saturating concentration of a red membrane impermeant Halo ligand (JF633i) followed by a pulse of green membrane permeant Halo ligand (JF585) to show that when expressed in neurons, this NaCT is distributed between intracellular and PM pools at both the cell soma and in axons (Figure S3 e). To quantitatively assess the distribution of overexpressed transporter and to gain insight into the intracellular compartment, we used a membrane permeable and pH sensitive HaloTag ligand, Virginia Orange (VO)^35^ (Figure 3 i). A brief superfusion of an extracellular acidic buffer led to strong and rapid quenching in a portion of the fluorescence signal, while superfusion of NH_4_Cl, which will alkalize any internal acidic compartment ^36^, led to an increase in VO fluorescence (Fig. 3 j). Making use of the known pKa of Halo-bound VO allowed us to determine that the C-terminal tagged SLC13A5 has a surface fraction of ∼ 50% (Fig. 3 k) while the remainder resides in an intracellular compartment with a pH ∼ 6.7 (Fig. 3 l).

Using the same C-terminal HaloTag construct and VO ligand, we were able to demonstrate that the intracellular compartment containing SLC13A5 is not being exocytosed in response to AP firing (Figure S3 f). Instead, we found that the compartment is slowly recycled under steady state, which is evident from replacement of surface SLC13A5-Halo stained with membrane impermeable dye of one color by a SLC13A5-Halo with membrane impermeable dye of another color at a rate of approx. 4% of total SLC13A5 in 20 minutes (Figure S3 g, j).

Overall, our results show that in neuronal soma, SLC13A5 is localized at TGN, while in the axons, it is localized on a slowly recycling intracellular vesicles with luminal pH of 6.7, and that the intracellular compartment is responsible for the control of glycolytic activation by electrical activity.

### Neuronal activity results in SLC13A5 dependent clearance of axonal citrate

These data all predict that during electrical activity the concentration of cytoplasmic citrate would diminish through the action of SLC13A5, presumably by sequestering into an intracellular endosomal pool of vesicles. To test this hypothesis, we used Citroff1 to examine the dynamics of cytoplasmic citrate in axons during and after a burst of AP firing. Our superfusion citrate experiments (Fig. 3 a-d) showed that resting cytoplasmic citrate levels are likely much lower than the apparent K_D_ of the sensor, and therefore it is imperative to properly control for possible artifacts since one will be dealing with relatively small signal changes. To this end we deployed the citrate-insensitive variant of Citroff1, which reported small AP triggered increase in apparent citrate (Fig. 4 a) (this is actually a decrease in fluorescence signal, but as Citroff1 is an inverse sensor, we are reporting the 1/F signal, see methods). This apparent increase in citrate very likely arises from a transient acidification of the axonal cytoplasm driven by the burst of AP firing^17^. In contrast, Citroff1 showed a drop in citrate signal, initially partially masked by the likely pH effect (Fig. 4 a) that reached a minimum ∼ 15 s after the end of the stimulus train and slowly recovered over the next ∼ 30 s and overshot the initial resting value. The activity-driven drop in citrate was eliminated in SLC13A5 KD axons, while the recovery was not affected (Figure 3 b, c), suggesting that SLC13A5 is responsible for activity dependent drop of axonal citrate. The magnitude of this citrate drop was not altered when axons were additionally superfused with extracellular citrate (Figure S4 a, b) as in Figure 3, consistent with the idea that the flux of citrate we measured during electrical activity is not influenced by the gradient in citrate concentration between the cytoplasm and the extracellular space, as would have been expected if this was a plasma membrane NaCT responsible for citrate clearance.

**Figure 4.**
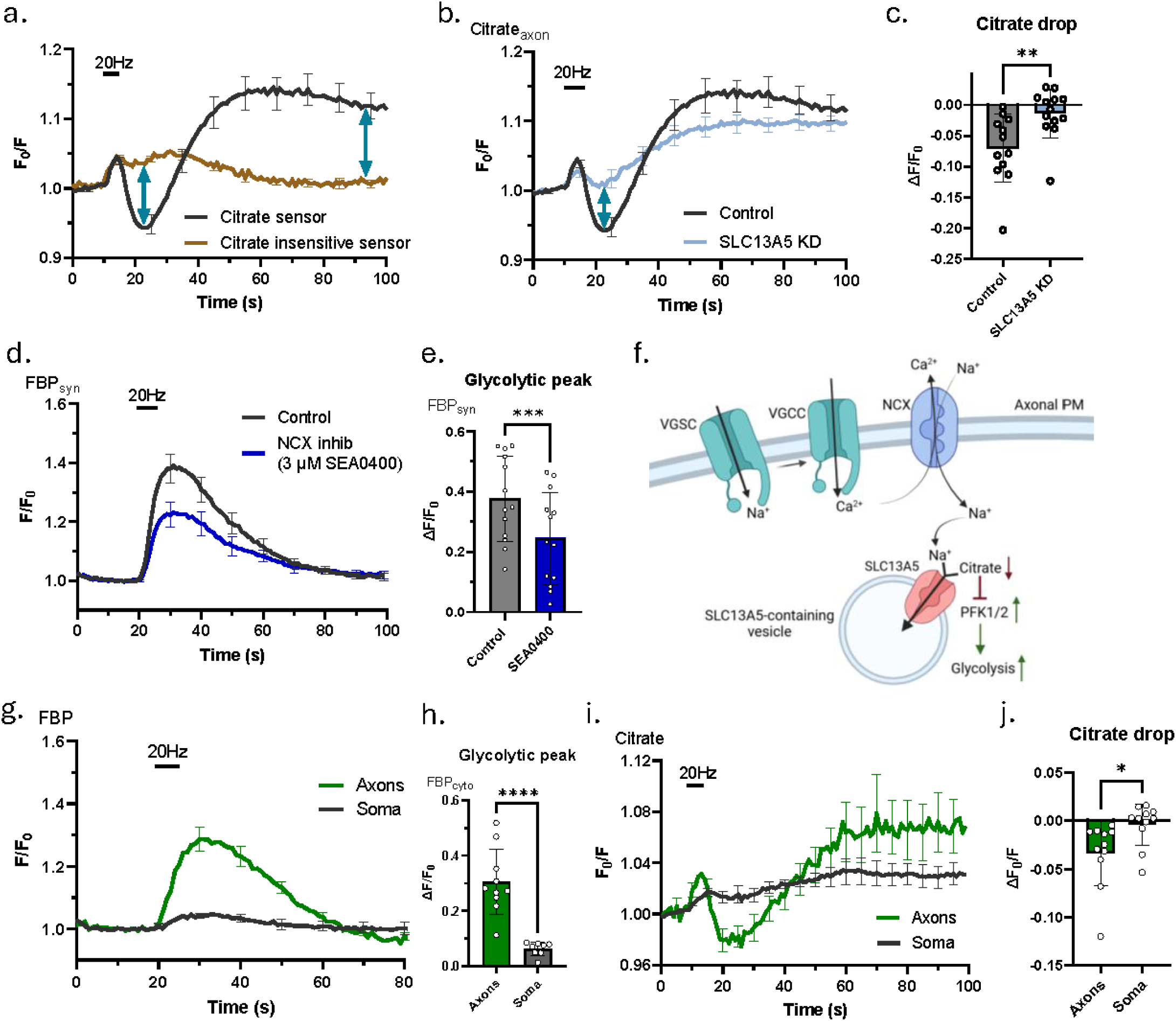
Neuronal activity results in SLC13A5 dependent clearance of axonal citrate. (a) Average +/-SEM axonal Citroff1 (black) and CitroffRH (brown) traces in response to 100 APs at 20 Hz. (b) Average +/-SEM axonal Citroff1 traces of control (black) and SLC13A5 KD (light blue) neurons in response to 100 APs at 20 Hz. (c) Mean citrate drop (lowest Citroff1 signal drop below baseline) +/-SEM in response to 100 APs at 20 Hz, quantified for control (n=12) and SLC13A5 KD (n=13). (d) Average +/-SEM synapto-Hylight traces for control (black) and SEA0400 treated (dark blue) neurons stimulated with 100 APs at 20 Hz. (e) Mean glycolytic peak +/-SEM in response to 100 APs at 20 Hz, quantified for control (n=13) and SEA0400 treated (n=13) neurons. Paired t-test. (f) Schematic illustrations of proposed mechanism for SLC13A5 dependent control of presynaptic glycolysis. VGSC – voltage gated sodium channel, VGCC – voltage gated calcium channel. (g) Average +/-SEM cytosolic Hylight traces of neuronal somas (black) and axons (green) stimulated with 100 APs at 20 Hz. (h) Mean glycolytic peak +/-SEM in response to 100 APs at 20 Hz, quantified for neuronal somas (n=8) and axons (n=10). (i) Average +/-SEM axonal Citroff1 traces of neuronal somas (black) and axons (green) stimulated with 100 APs at 20 Hz. (j) Mean citrate drop +/-SEM in response to 100 APs at 20 Hz, quantified for neuronal somas (n=11) and axons (n=11). Paired t-test was used for (e) and (j).

SLC13A5 is a Na^+^-dependent citrate transporter and, as the intracellular concentration of Na^+^ ([Na^+^]) is much lower than the extracellular [Na^+^], we wondered where the driving force for citrate extrusion into an intracellular compartment could arise. During electrical activity synaptic terminals take up extracellular Ca^2+^ which in turn controls many aspects of neurotransmitter release^37^. This Ca^2+^ is extruded across the plasma membrane via the activity of both a Ca^2+^ ATPase (PMCA), and a sodium-calcium exchanger (NCX)^38,39^.The latter exchanges three Na^+^ ions for each extruded Ca^2+^ ion and creates a transient Na^+^ load in the cytoplasm. We reasoned that that Na^+^ entering via NCX might provide the driving force for SLC13A5 acting on an intracellular pool of vesicles. Application of a known NCX inhibitor, SEA0400^40^ resulted in blunted activation of presynaptic glycolysis (Figure 4 d, e), suggesting that NCX activity likely serves as a driving force for Na^+^/citrate co-transporter (Figure 4 f).

The mechanism we described here provides an explanation of SLC13A5 dependent metabolic activation at presynaptic terminals and axons. We also evaluated activity dependent glycolysis in neuronal cell bodies to provide a comparison with our measurements in axons. Unlike in the axon, bursts of AP firing led only to a small increase of the FBP signal in cell bodies (Figure 4 g, h). We then examined the dynamics of cytoplasmic citrate in the cell soma, to determine if the suppressed glycolytic activation might be due to a lower citrate clearance driven by electrical activity in this cellular region. These experiments demonstrated that activity-driven citrate clearance is likely restricted to axons (Figure 4 i, j). These results emphasize preferential importance of SLC13A5 dependent mechanism of glycolytic activation for presynaptic terminals, a claim that is further supported by a larger ATP deficit in presynaptic boutons compared to somas in SLC13A5 KD neurons (Figure S4 c). Given that citrate serves as an inhibitor of PFK1 and, as a result, of glycolysis, this transient SLC13A5 dependent drop of citrate level is likely responsible for glycolytic activation during action potential firing and represents a novel form of axonal metabolic control.

### Loss of SLC13A5 degrades synaptic function that can be reversed by a therapeutic activator of glycolysis

We addressed the physiological consequences of loss of SLC13A5 on synaptic function using a well-established technique based on synaptic vesicle (SV) targeted pH probe, vGlut1-pHluorin^41^ (vG-pH) (Figure 5 a). SV recycling is acutely sensitive to impairment in bioenergetics and therefore serves as a good metric for assessing the impact of genetic mutations in metabolism on synapse function^18^. Here we made use of a variant of our synaptic endurance test^16^, whereby neurons are subject to repeated bouts of activity at minute intervals and vG-pH signals are tracked to determine the efficacy of SV recycling for each stimulus round. In control neurons, SV recycling was well sustained for 12 rounds of repetitive AP trains, while SLC13A5 KD neurons failed to fully recover after the 5th stimulus burst (Figure 5 b, c). This failure to recycle SVs is typically indicative of defective ATP production, as was shown before^17^ and is an expected consequence of inadequate glycolytic activation. This inability to sustain presynaptic endurance was rescued upon reintroduction of the C-terminally tagged, but not N-terminally tagged human SLC13A5, once again emphasizing the importance of proper localization of the transporter for its activity dependent function (Figure S5 a).

**Figure 5.**
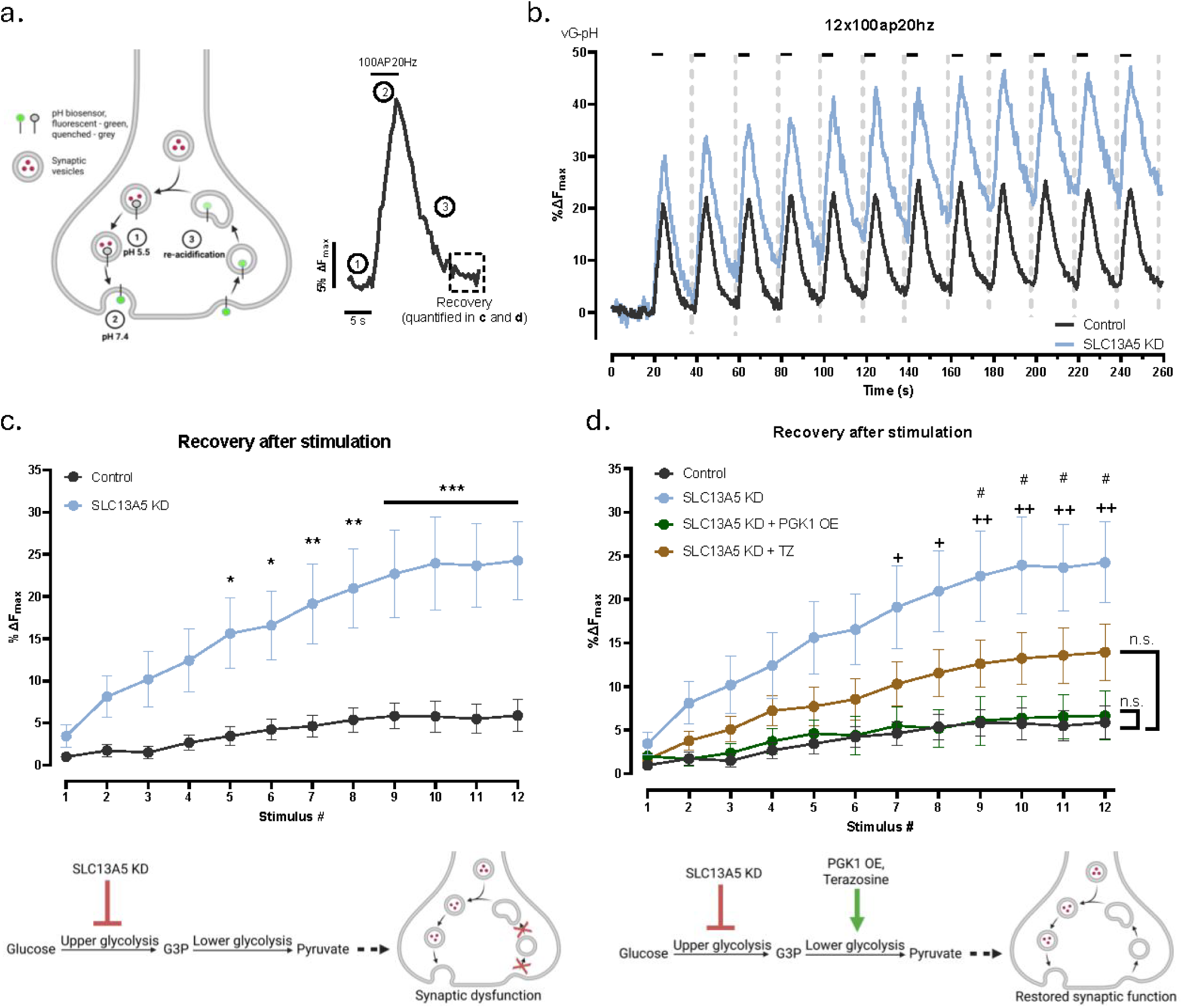
Loss of SLC13A5 degrades synaptic function that can be reversed by a therapeutic activator of glycolysis. (a) Schematic illustration of vG-pH based synaptic vesicle recycling assay (left) with a corresponding representative vG-pH trace in response to a train of 100 APs at 20 Hz (right). (b) Average vG-pH trace of control (black) and SLC13A5 KD (light blue) neurons in response to 12 trains of 100 APs at 20 Hz. Dashed lines show areas used for analysis in (c). (c and d) Recovery after stimulation, quantified as remaining vG-pH fluorescence following AP train, for (c) control (black, n=19) and SLC13A5 KD (light blue, n=14) neurons, (d) with addition of SLC13A5 KD + PGK1 OE (green, n=7) and SLC13A5 KD + TZ treated (brown, n=17) neurons. Two-way ANOVA with (c) Šídák’s and (d) Tukey’s multiple comparisons test. Lower panels schematically illustrate mechanism of synaptic dysfunction caused by SLC13A5 deficiency (c) and the proposed therapeutic intervention (d).

Recently we demonstrated that during action potential firing, phosphoglycerate kinase 1 (PGK1) is the rate limiting enzyme in nerve terminal glycolysis, and modest changes in the concentration of this enzyme in nerve terminals can protect synaptic function against hypometabolism *in vitro* and *in vivo*^16^. As PGK1 is down stream of PFK1 in the glycolytic cascade, we reasoned that a deficit in PFK1 function could potentially be rescued by increased PGK1 function, since the latter is rate-limiting. Overexpression of PGK1 in SLC13A5 KD neurons completely reversed the synapse dysfunction in the synaptic endurance assay (Figure 5 d). Our previous work on PGK1 function at nerve terminals was motivated in part by the discovery that this enzyme is activated as an off-target effect by the FDA approved treatment for benign prostate hyperplasia (BSH), Terazosin (TZ) and that TZ was shown in retrospective analysis to significantly lower the risk of Parkinson’s Disease (PD)^42,43^. Our own experiments previously demonstrated that TZ application in primary neurons leads to a dramatic acceleration of presynaptic ATP production, about ∼ half of that achieved with a greater than 2-fold overexpression of PGK1^16^. Similar to PGK1 overexpression, treatment of SLC13A5 KD neurons with TZ led to a strong suppression of the synapse dysfunction in the synaptic endurance assay (Figure 6 d). The fact that alternate glycolytic activators that act downstream of PFK1 can suppress the physiological impairment in nerve terminal function driven by loss of SLC13A5 strongly supports our mechanistic interpretation that this NaCT deficiency leads to an axonal bioenergetic deficit. Furthermore, these results provide a promising therapeutic strategy to alleviate the consequences of SLC13A5 deficiency.

## Discussion

The brain imposes a significant metabolic burden on the body, requiring constant substrate delivery to sustain neural activity and their complex sequalae. Consequently, supply and use of glucose, the main energy substrate of the brain, needs to be well managed. One of the possible adaptations to such constant energy demand while substrate is limiting is an efficient gating mechanism that would allow on-demand energy production. Our work shows that neurons suppress their glycolysis through metabolic feedback loop involving PFK1 and citrate (Figure 2). This might be an adaptive mechanism to conserve glucose and keep glycolysis low at resting state, while neuronal activity temporarily relieves this inhibition through SLC13A5 dependent axonal citrate clearance (Figure 4 a-c). Thus, this NaCT based gating links energy production to neuronal AP firing, providing increased metabolic bandwidth when it is most needed (Figure 1 d). The inability to properly boost metabolism during activity is a likely underlying cause of the neurological manifestation of SLC13A5 mutations. We speculate that EOEE might result from increased susceptibility of certain types of neurons to activity dependent bioenergetic deficits, leading to synaptic dysfunction and unbalanced brain circuits.

Besides the brain, mammalian SLC13A5 is expressed in liver and testis^44^. Based on citrate uptake assays in hepatocytes^13^, NaCT is likely expressed on the PM in liver cells. Thus, SLC13A5 dependent metabolic control mechanism described here for neurons is likely different than in the liver, as it mediates uptake of extracellular citrate in this tissue, where it is thought to function in controlling lipid synthesis^45^. In neurons its function in sequestration of intracellular citrate not only requires localization to an internal organelle (Figure 3), but also would require changes in intracellular Na^+^ to drive transient changes in activity (Figure 4 d-f). Thus, the glycolytic gating function of SLC13A5 would only operate in excitable cells. We speculate that cell type specific differential localization of the NaCT might shed light into tissue specific phenotypes of SLC13A5 deficiencies in different species. SLC13A5 in cell types where it is expressed on the PM will likely function in ways that are unrelated to membrane excitability, since the Na^+^ gradient will always be large and in a direction that favors citrate uptake, which in turn can impact lipid synthesis. Mutations in SLC13A5 in these tissues might therefore provide protective effects in high fat diet^14^. In neurons, in contrast, where the NaCT functions on internal membranes, mutations impact glycolytic gating that have deleterious outcomes. Our work provides an important cell-biological link between protein localization and function that is clinically relevant. Further studies focusing on identity of the slow-recycling NaCT carrying compartment and cell-type specific targeting mechanism of the SLC13A5 are needed.

Detailed mechanistic insight into metabolic mechanism of action of the NaCT in neurons allowed us to devise a therapeutic approach to combat synaptic dysfunction in SLC13A5 deficient neurons. Since neurons lacking SLC13A5 have deficient activity dependent glycolysis (Figure 2), it would theoretically mimic neuronal activity in hypometabolic conditions. While overexpression of PFK1 didn’t restore activity dependent glycolysis (Figure 2 b, f), genetic and pharmacological upregulation of another critical glycolytic enzyme, PGK1, which we previously showed is the rate-limiting enzyme in ATP production at nerve terminals during bursts of activity^16^, rescued neurons from synaptic dysfunction caused by SLC13A5 KD (Figure 5 d). Reversal of synaptic dysfunction by Terazosin is particularly promising as it is an FDA approved drug, which makes further studies using TZ in SLC13A5 deficiency models and patients appealing.

In summary, this work provides a mechanistic insight into a novel metabolic gating of presynaptic energy metabolism, improves our understanding of a genetic disorder, and provides a potential therapeutic strategy to combat it. In light of the novelty, the study has certain shortcomings that need to be addressed in future work, including the identity of relevant intracellular compartments, development of citrate biosensors with fitting dynamic range, and testing of TZ on animal models with EOEE.

### Contact for Reagent and Resource Sharing

Further information and requests for resources and reagents should be directed to and will be fulfilled by the Lead Contact, Timothy A. Ryan at.

## Acknowledgements

We thank members of the Ryan laboratory for their valuable suggestions and input on this work. This work was supported in part by the NIH (TAR: NS036942 and NS11739) and the Simons Foundation (Z.K.). The graphical abstract and schematic illustrations in Figures 2-5, S3 were created using Biorender.com.

## Author Contributions

Conceptualization, Z.K. and T.A.R.; methodology, Z.K., A.C.K. and T.A.R.; Investigation, Z.K. (all experiments, except for high-resolution Airy scan imaging), L.B. (high-resolution Airyscan imaging); writing—original draft, Z.K. and T.A.R.; writing—review & editing, all co-authors; funding acquisition, Z.K. and T.A.R.; resources, T.A.R. and J.L.S.; supervision, T.A.R.

## Supplementary data

**Figure S1.**
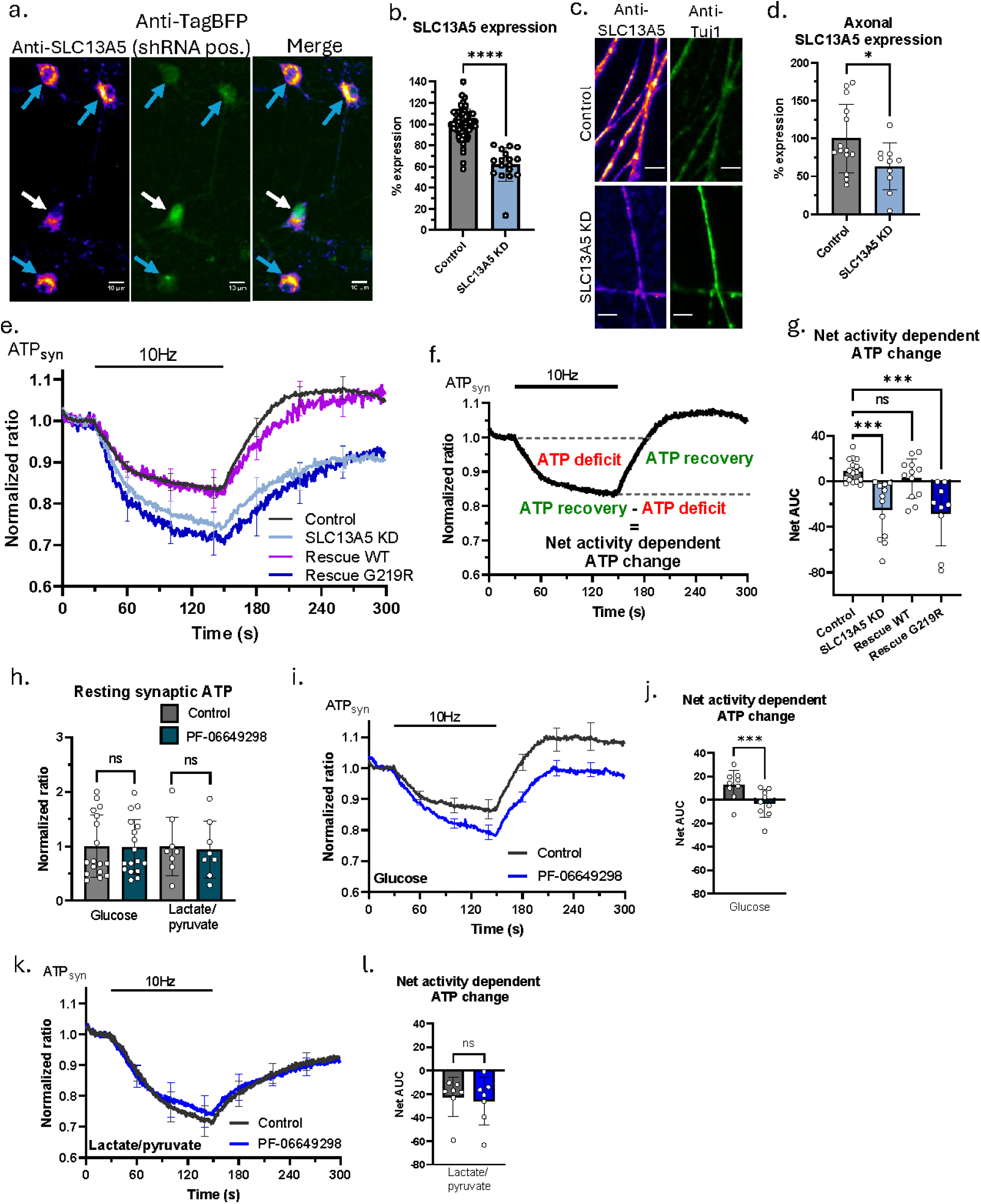
SLC13A5 is needed for activity dependent ATP production from glucose. (a) Representative immunofluorescence images of control (blue arrows) and SLC13A5 KD (white arrow) hippocampal neurons stained with anti-SLC13A5 antibody (pseudo-color) and anti-TagBFP nanobody (green). Scale bar is 10 µm. (b) Mean SLC13A5 expression +/-SEM in somas of control (n=65) and SLC13A5 KD (n=17) neurons. (c) Representative immunofluorescence images of axons from control (upper) and SLC13A5 KD (lower) hippocampal neurons stained with anti-SLC13A5 (pseudo-color) and anti-Tuj1 (green) antibodies. (d) Mean axonal SLC13A5 expression +/-SEM in control (n=14) and SLC13A5 KD (n=10) neurons. (e) Average +/-SEM synapto-iATPSnFR2-miRFP670nano3 traces for control (black), SLC13A5 KD (light blue), rescue with WT SLC13A5 (magenta), and rescue with G219R mutant SLC13A5 (dark blue) neurons stimulated with 1200 APs at 10 Hz. (f) Average control synapto-iATPSnFR2-miRFP670nano3 trace with markings showing AUC used for quantification of the net activity dependent ATP change. (g) Mean presynaptic “net activity dependent ATP change” +/-SEM, quantified as shown in (f) for control (n=17), SLC13A5 KD (n=12), Rescue with WT SLC13A5 (n=12), and Rescue with G219R mutant of SLC13A5 (n=9) neurons. (h) Mean resting synaptic ATP +/-SEM depicted as control-normalized iATPSnFR2-miRFP670nano3 ratio for control with glucose (n=17), PF-06649298 with glucose (n=17), control with lactate/pyruvate (n=8), and PF-06649298 with lactate/pyruvate (n=8). (i) Average +/-SEM synapto-iATPSnFR2-miRFP670nano3 traces for control (black) and PF-06649298 treated (blue) neurons stimulated with 1200 APs at 10 Hz in glucose containing buffer. (j) Mean net activity dependent ATP change +/-SEM, for control (n=9) and PF-06649298 treated (n=9) neurons in glucose containing buffer. (k) Average +/-SEM synapto-iATPSnFR2-miRFP670nano3 traces for control (black) and PF-06649298 treated (blue) neurons stimulated with 1200 APs at 10 Hz in lactate/pyruvate containing buffer. (l) Mean net activity dependent ATP change +/-SEM, for control (n=7) and PF-06649298 treated (n=7) neurons in lactate/pyruvate containing buffer.

**Figure S2.**
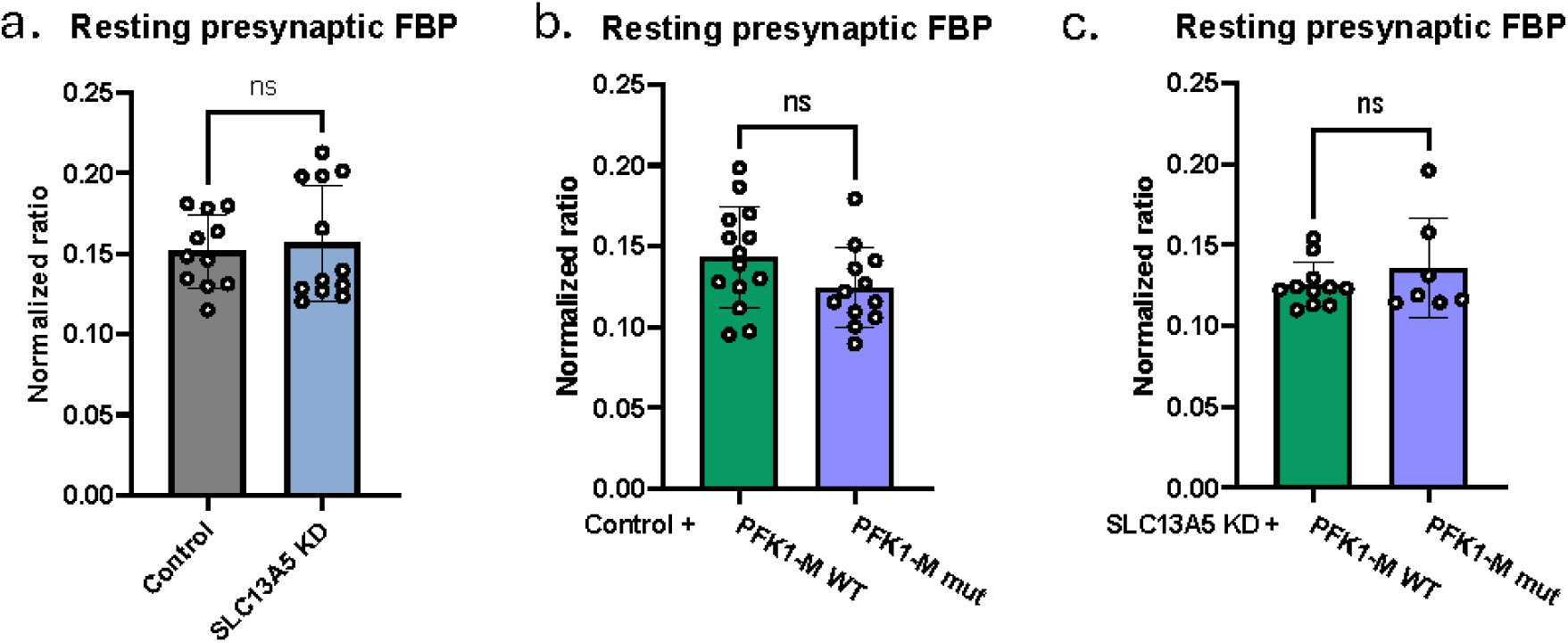
SLC13A5 KD and PFK1 OE don’t affect resting presynaptic FBP levels. (a-c) Mean resting presynaptic FBP +/-SEM, quantified as Hylight green-to-blue excitation ratio, for (a) control (n=11) and SLC13A5 KD (n=12), (b) control PFK1-M WT OE (n=14) and control PFK1-M mut OE (n=12), (c) SLC13A5 KD PFK1-M WT OE (n=11) and SLC13A5 PFK1-M mut OE (n=7) neurons.

**Figure S3.**
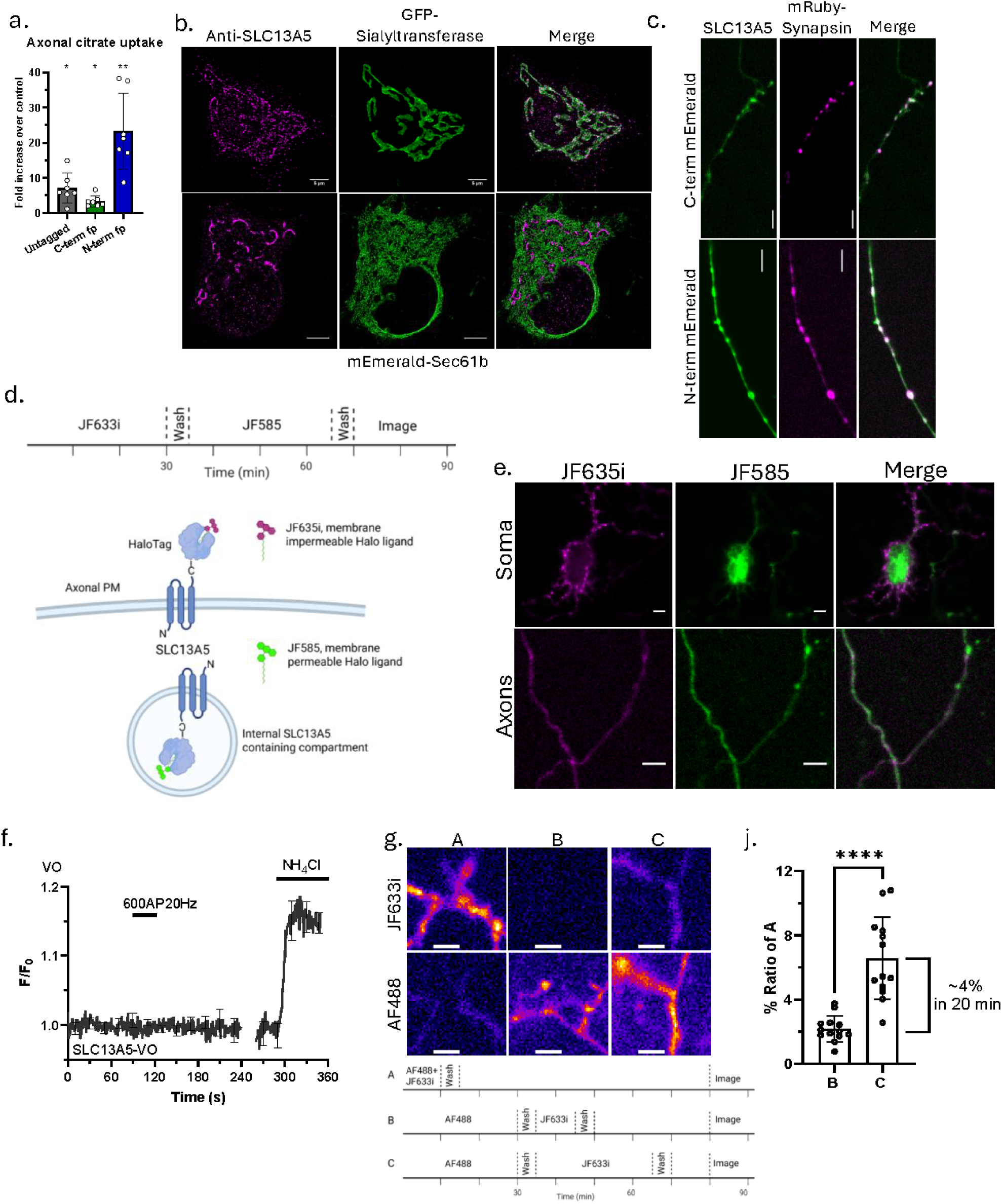
Intracellular SLC13A5 localization. (a) Mean fold increase in axonal citrate uptake over control +/-SEM in response to 1 mM citrate superfusion, quantified for neurons overexpressing untagged (n=7), c-terminally tagged (n=7), and n-terminally tagged (n=7) SLC13A5 fusion constructs. (b) Representative high-resolution Airyscan images of neuronal soma showing endogenous SLC13A5 staining (both left panels), GFP-Sialyltransferase (middle upper panel), mEmerald-Sec61b (middle lower panel) and merged images (right panels). Scale bar is 5 µm. (c) Representative images of neurons expressing c-, and n-terminally mEmerald-tagged SLC13A5 fusion proteins co-transfected with mRuby-Synapsin. Scale bar is 5 µm. (d) Experimental protocol (upper panel) and schematic illustration of predicted intracellular localization of SLC13A5-Halo construct and labelling with membrane impermeable (JF635i) and permeable (JF585) Halo-Tag ligands (lower panel). (e) Representative images of neurons expressing SLC13A5-Halo construct and labelled according to the protocol illustrated in (d). Images show PM and intracellular localization of SLC13A5-Halo construct. (f) Average +/-SEM VO trace in response to 600AP at 20 Hz and NH_4_Cl superfusion. (g) Representative images of neurons expressing SLC13A5-Halo construct labelled with membrane impermeable Halo-Tag dyes (JF635i and AF488) according to the protocol shown below. Scale bar is 4 µm. (j) Analysis of the experiment shown in (g); JF635i to AF488 ratios of experimental conditions B and C were normalized as percentages of A, and reported as mean +/-SEM, n=13. The difference between ratios of C and B is 4%, while incubation time difference is 20 minutes, corresponding to 4% recycling in 20 minutes.

**Figure S4.**
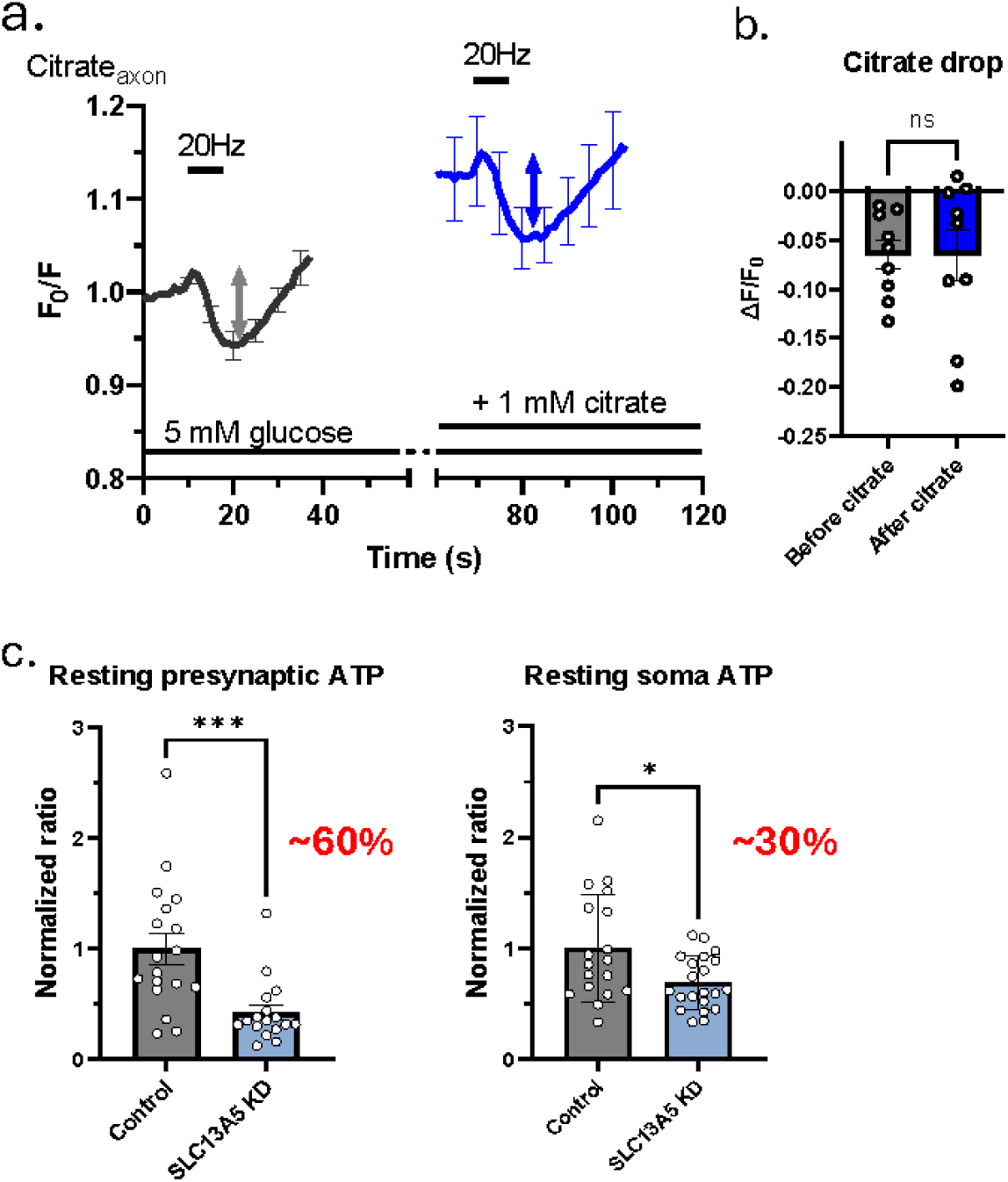
SLC13A5 preferentially functions in axons. (a) Average +/-SEM axonal Citroff1 traces of control neurons before (black) and after 1 mM citrate superfusion (blue) in response to 100 APs at 20 Hz. (b) Mean citrate drop +/-SEM in response to 100 APs at 20 Hz, quantified for neurons before (n=9) and after 1 mM citrate superfusion (n=9). (c) Mean resting presynaptic (left) and soma (right) ATP +/-SEM, depicted as iATPSnFR2-miRFP670nano3 ratio, normalized to respective controls. Control presynaptic, n=18; SLC13A5 KD presynaptic, n=17; Control soma, n=18; SLC13A5 KD soma, n=22.

**Figure S5.**
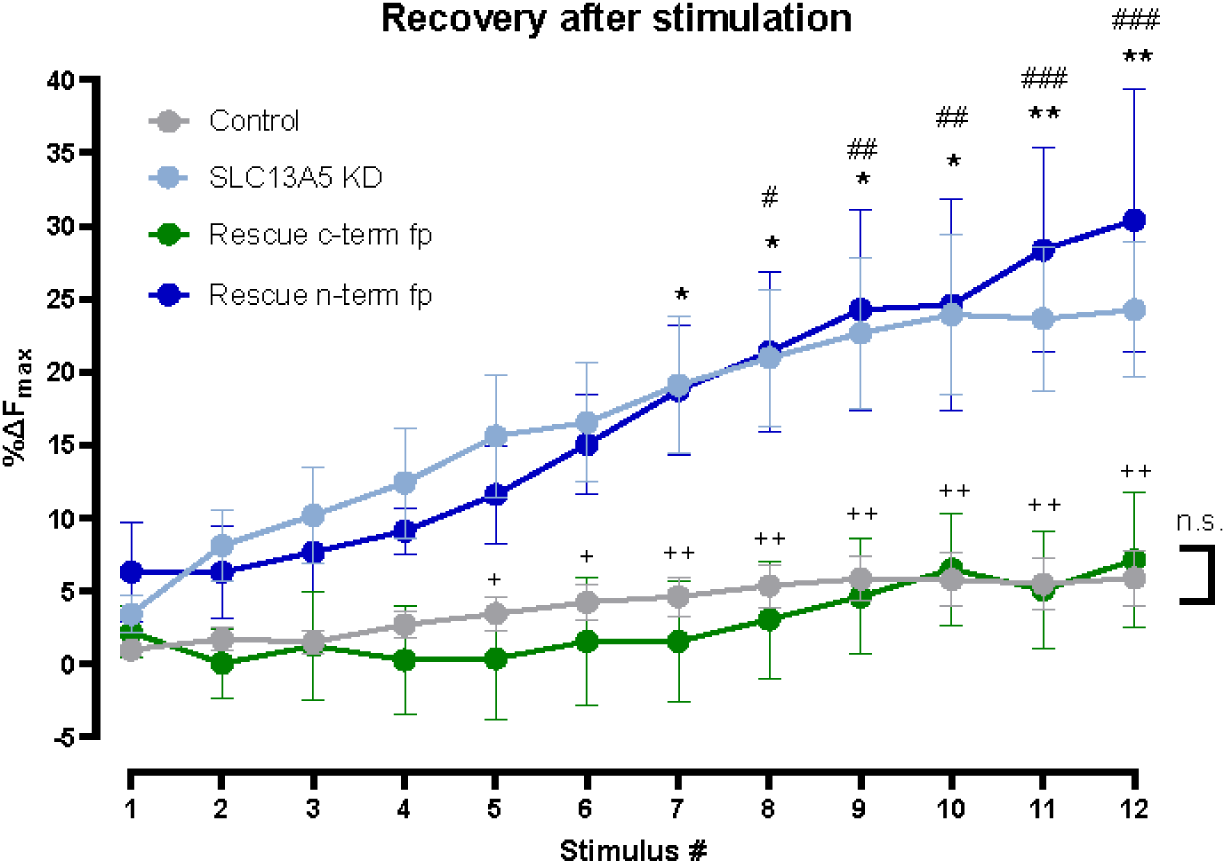
Proper SLC13A5 localization matters for synaptic function. Recovery after stimulation, quantified as remaining vG-pH fluorescence following AP train, for control (black, n=19), SLC13A5 KD (light blue, n=14), Rescue with c-terminally tagged SLC13A5 (green, n=7), and rescue with n-terminally tagged SLC13A5 (brown, n=17) neurons.

## STAR METHODS

### KEY RESOURCES TABLE

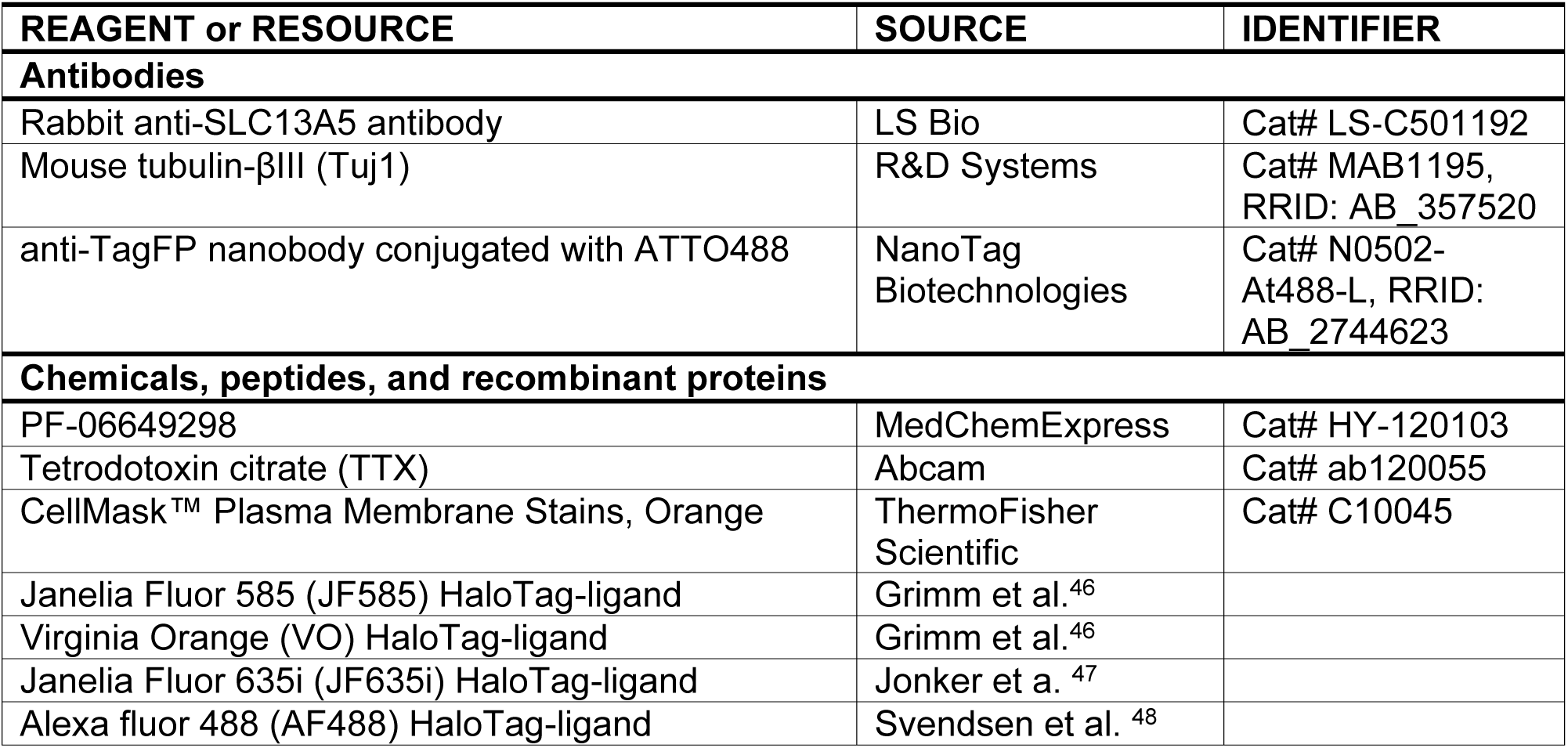
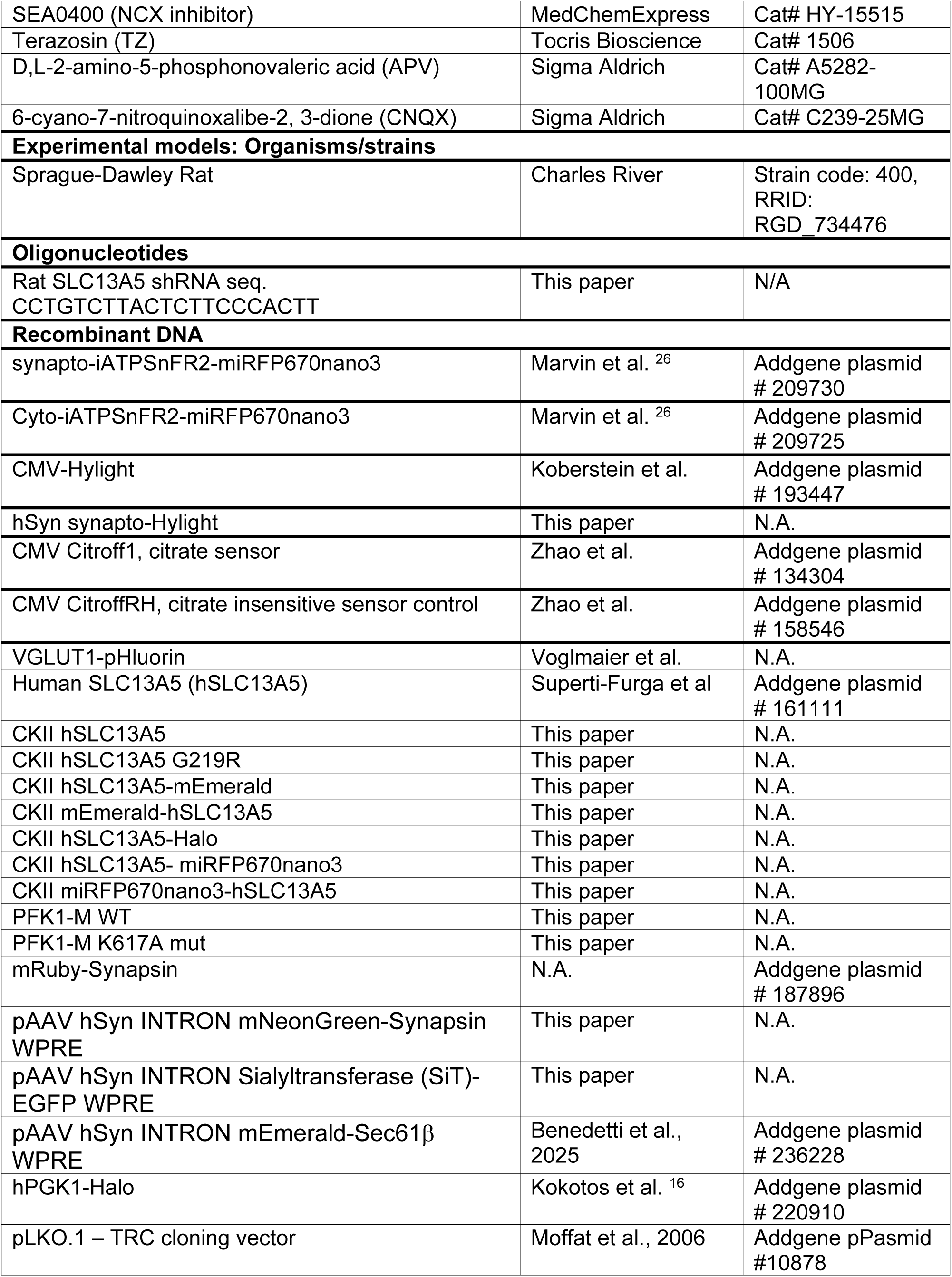
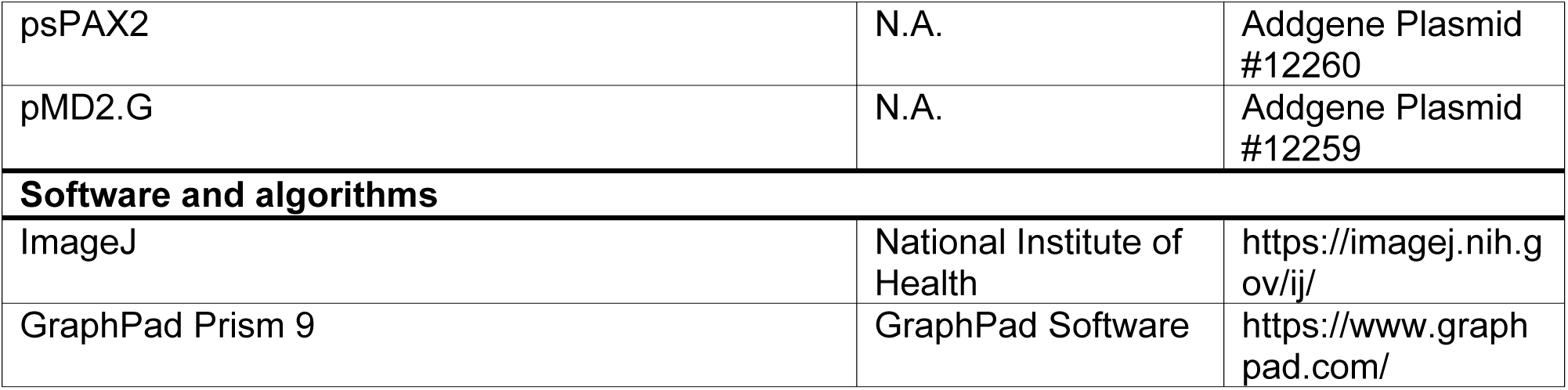

### EXPERIMENTAL MODEL AND STUDY PARTICIPANT DETAILS

#### Animals

All animal-related experiments were performed in accordance with protocols approved by the Weill Cornell Medicine IACUC, protocols number 0601-450A and 2009-0026 and Janelia Research Campus IACUC, protocol number 21-206. Wild-type rats were of the Sprague-Dawley strain (Charles River Laboratories, strain code: 400, RRID: RGD_734476).

#### Primary neuronal culture

Hippocampal CA1-CA3 neurons were isolated from 1- to 2-day-old rats of mixed gender, plated on poly-ornithine-coated coverslips, transfected 7 days after plating, and imaged 14-21 days after plating as previously described (dx.doi.org/10.17504/protocols.io.ewov1qxr2gr2/v1). For Airyscan Microscopy experiments, primary cultures of hippocampal neurons were prepared from rat brains of both genders at embryonic day 19 (E19). The hippocampi were dissected in cold Hank’s Balanced Salt Solution (Sigma, H2387), which was supplemented with 1 mM HEPES (Fisher, BP310), 4 mM NaHCO3 (Sigma-Aldrich S5761), and 20% FBS (Biotechne, S11550H). The cells were then dissociated by tissue trituration and papain treatment (Worthington Biochemical Corporation, LK003178, PDS Kit) in HBSS at pH 7.4 at 37°C for 10 minutes. Live, dissociated cells (Trypan Blue exclusion) were counted and plated on 25 mm round coverslips (Electron Microscopy Science, 72196-25) coated with Poly-L-Ornithine Solution (0.01%, Sigma-Aldrich, P4957) and 10 μg/ml laminin (Sigma-Aldrich, 11243217001) in DPBS (Thermofisher, 14040182) at a density of 3.4 x 104 cells/cm2 in NeuroCultTM Neuronal Plating Medium (STEMCELL Technologies, 05713) supplemented with 2% NeuroCultTM SM1 neuronal supplement (STEMCELL Technologies, 05711) and 2 mM GlutaMAX™ Supplement (Gibco, 35050061). Five days after plating, half of the plating medium was removed from each well and replenished with the same volume of BrainPhys™ Neuronal Medium (STEMCELL Technologies, 05792) supplemented with 2% NeuroCult™ SM1 Neuronal supplement. The cells were maintained in vitro at 37°C and 5% CO2. From DIV 5 onwards, the medium was refreshed every four days by replacing half of the medium with BrainPhys™ neuronal medium supplemented with 2% NeuroCult™ SM1 neuronal supplement.

### METHOD DETAILS

#### Plasmids

The following previously published DNA constructs were used: synapto-iATPSnFR2-miRFP670nano3^26^, cyto-iATPSnFR2-miRFP670nano3^26^, Hylight^29^, Citroff1^32^, CitroffRH (citrate insensitive control of Citroff1)^32^, VGLUT1-pHluorin^41^, hPGK1-Halo^16^. Short-hairpin RNA (shRNA) sequence against SLC13A5 was designed using GPP Web Portal (https://portals.broadinstitute.org/gpp/public/). The shRNA oligonucleotides were cloned into pLKO-TRC vector and expressed under U6 promoter. The pLKO-TRC plasmid also expresses nuclear targeted TagBFP under hPGK promoter as a selection marker. Synapto-Hylight was generated by sub-cloning Hylight into a vector with hSynapsin promoter and synaptophysin targeting sequence. Human SLC13A5 construct was a gift from RESOLUTE Consortium & Giulio Superti-Furga (Addgene plasmid # 161111; http://n2t.net/addgene:161111; RRID:Addgene_161111). SLC13A5 was subcloned under CKII promoter. Point mutation was generated by PCR cloning. SLC13A5 fusion constructs were generated by n-, or c-terminally subcloning miRFP670nano3, mEmerald, or Halo-tag with a short flexible linker. Human PFK1-M K617A point mutation was generated by PCR from WT human PFK1-M.

pAAV hSyn INTRON mNeonGreen-Synapsin WPRE and pAAV hSyn INTRON Sialyl transferase (SiT)-EGFP WPRE were generated by subcloning the cORF of mNeonGreen-Synapsin (from the Ryan lab) and SiT-GFP (a gift from Jennifer Lippincott-Schwartz, Addgene #166943, from Patterson GH et al., Cell 2008), respectively, at BamHI and HindIII sites of pAAV Syn intron Psam4GlyR m3m4 HA (a gift from Scott Sternson, Addgene #196041). PCR amplification of the fragments and their subsequent ligation was performed using the In-Fusion^®^ HD Cloning Kit and online tools (BD Clontech, Takara Bio, USA. pAAV hSyn INTRON mEmerald-Sec61β WPRE is a gift from Jennifer Lippincott-Schwartz, Addgene # 236228 (from Benedetti et al., Cell 2025). All plasmids were verified by sequencing with Plasmidsaurus, (Eugene, OR, USA).

#### shRNA lentivirus generation

HEK 293FT cells were transfected with the SLC13A5 shRNA carrying pLKO plasmid along with two packaging plasmids (pPAX2 and pMD2) using JetPRIME DNA/siRNA transfection reagent (Polyplus, 101000027). Transfection media was replaced with virus production media 16 hours after transfection. The supernatant containing lentivirus was collected at 46-48 hours, filtered through 0.45 μm syringe filter set and concentrated using Lenti-X concentrator (Takara Bio, 631231). The virus titer was measured using Lenti-X GoStix Plus assay kit (Takara Bio, 631280). Viral KD of SLC13A5 was used for KD verification in neuronal axons.

#### AAV particle production

The Janelia Viral Tools team at HHMI Janelia Research Campus (Ashburn, VA) produced adeno-associated virus (AAV). AAV particles were added to DIV5 neurons, and media exchange was performed 2-4 days after infection. Primary neurons were fixed and processed for immunofluorescence after 10 days of viral transduction.

#### Live-cell imaging

Live-cell imaging was performed as previously described dx.doi.org/10.17504/protocols.io.q26g7pn4qgwz/v1). All imaging experiments were performed on a custom-built laser illuminated epifluorescence Zeiss Axiovert 200 microscope with Andor iXon camera (model: DU-897U-CS0-BVF) and 40X 1.3 NA Fluar Zeiss objective. Cells were maintained at 37 ^0^C and constantly perfused at a rate of 0.1 ml/min with a Tyrode’s solution containing 5 mM glucose (1 mM glucose for VGLUT1-pH experiment), 119 mM NaCl, 2.5 mM KCl, 2 mM CaCl2, 2 mM MgCl2, 50 mM HEPES (pH 7.4). The buffer was supplemented with 0.01 mM 6-cyano-7-nitroquinoxalibe-2, 3-dione (CNQX) and 0.05 mM D,L-2-amino-5-phosphonovaleric acid (APV) to suppress post-synaptic responses.

Action potentials were evoked by passing 1-ms current pulses, yielding fields of approximately 10 V cm−1 via platinum-iridium electrodes. A385 High Current Stimulus Isolator from World precision Instruments was used.

Cell Mask Orange PM stain was used in 1:2000 dilution for 5 minutes at 37°C and 5% CO_2_ in Tyrode’s solution. Following incubation, cells were washed 3 times with Tyrode’s solution and imaged.

For all imaging experiments, background-subtracted images were used for further analysis.

#### ATP measurements

For synapto-iATPSnFR2-miRFP670nano3 experiments, nerve terminals were selected in the miRFP670nano3 channels, blind to the iATPSnFR2 channel. Steady-state levels and dynamic traces are reported as iATPSnFR2 (green) to miRFP670nano3 (far-red) ratio, normalized to control ratio.

For TTX treatment experiments, 500 nM TTX was added 48-72 hours before measurement in culture media. For experiments after TTX wash, TTX was washed with Tyrode’s buffer under constant superfusion for at least 10 minutes. PF-06649298 treatment was done in superfusion for 10 minutes before the measurement. For experiments with lactate and pyruvate, glucose containing Tyrode’s solution was replaced by glucose free, 1.25 mM lactate and 1.25 mM pyruvate containing Tyrode’s solution in superfusion 5 minutes before the measurement. Activity dependent ATP recovery was quantified as area under the curve (AUC) of synapto-iATPSnFR2-miRFP670nano3 traces after the end of AP train. The lowest point used for AUC calculation was the last point of AP train. Net activity dependent ATP change was quantified as AUC of the ATP trace starting from the last frame of electrical activity (also the lowest point), minus area above the curve (until normalized baseline) during and after electrical activity. Resting soma ATP was measured using untargeted, cytosolic iATPSnFR2-miRFP670nano3.

#### FBP measurements

For baseline blue/green ratiometric measurements, Hylight transfected neurons were illuminated with mercury arc lamp with custom filter sets. For blue light excitation and green emission, blue excitation filter, GFP dichroic and GFP emission filter was used. For green excitation and emission, green excitation filter, GFP dichroic and GFP emission filter was used. Dynamic measurements of FBP were done using 491 laser and GFP filter cube only. Baseline normalized traces were used for analysis. Glycolytic peak was quantified as the maximum increase of signal over baseline.

For NCX inhibitor experiments, 10 µM SEA0400 was applied in superfusion for 5-10 minutes before the measurement. For experiments comparing soma and axonal FBP dynamics, untargeted Hylight sensor was used.

#### Citrate measurements

Since Citroff1 is an inverse citrate sensor, baseline normalized fluorescence was inverted (1/F) for simplicity. For axonal citrate uptake experiments, maximal fluorescence change during 1 mM citrate superfusion was used for analysis. For activity dependent citrate drop evaluation, lowest Citroff1 signal drop below baseline after electrical stimulation was taken.

#### Halo-Tag based experiments

All Halo-tag ligands were used in 100 nM concentration and incubated with cells for 30 minutes at 37°C and 5% CO2 unless stated otherwise.

Pulse-chase labeling with membrane impermeable and permeable ligands was done as follows: SLC13A5-Halo transfected cells were incubated with membrane impermeable JF635i for 30 min, washed, incubated with membrane permeable JF585 for 30 min, washed, and imaged.

Experiments using Virginia Orange pH sensitive dye were done as follows: after VO labeling of SLC13A5-Halo transfected neurons, cells were perfused with Tyrode’s solution with short application of an acidic 5.5 pH MES buffered solution for 20 s to quench surface facing VO fluorescence; following pH recovery, cells were perfused with 7.4 pH 50 mM NH_4_Cl solution to alkalize internal VO. Surface fraction was determined as a fluorescence change evoked by short acid superfusion divided by maximum fluorescence after NH_4_Cl addition. Internal compartment pH was determined as previously reported^36^:

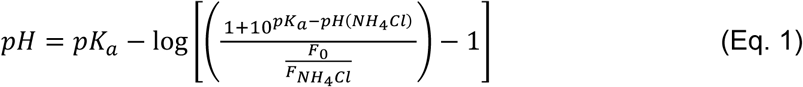

pK_a_ is the pK_a_ of VO, 6.46, pH(NH_4_Cl) is the pH of 50 mM NH_4_Cl buffer used, F_0_ is the fluorescence of VO measured before NH_4_Cl superfusion, F_NH4Cl_ is the fluorescence of VO measured upon NH_4_Cl superfusion when signal is stable.

For SCL13A5 recycling assay, SLC13A5-Halo transfected cells were: (A) labelled with both membrane impermeable dyes, JF635i and AF488, for 10 minutes, washed, and imaged; first labelled with AF488 for 30 minutes to saturate all extracellularly exposed SLC13A5-Halo, washed, and pulsed with JF635i for 10 (B) or 30 (C) minutes, washed, and imaged. The JF635i/AF488 ratio was quantified and normalized to JF635i/AF488 ratio of (A).

#### vGlut1-pH measurements

For synaptic vesicle recycling assay, the traces are reported as ΔF by subtracting the initial fluorescence signal before stimulation and normalized to total sensor fluorescence, revealed by the 50 mM NH4Cl Tyrode’s solution, yielding %ΔF_max_. Recovery after stimulation was quantified as average residual fluorescence before the start of the next AP train. For TZ treatment experiments, 10 µM TZ was added 24-48 hours before measurement in culture media.

#### Airyscan microscopy

Airyscan experiments were conducted on an inverted Zeiss LSM 980 microscope equipped with an Airyscan module. To excite different fluorescent proteins/dyes, lasers emitting 561 nm and 488 nm wavelengths were used; more specifically, AlexaFluorTM 568 using the 561 nm laser and mEmerald, mNeonGreen, or EGFP using the 488 nm laser. Imaging of neurons expressing nNeonGreen-Synapsin, (SiT)-EGFP or mEmerald-Sec61β after immunofluorescence staining with anti-SLC13A5 antibody were imaged in 15 DIV neurons with sequential detection of AlexaFluorTM 568 and mEmerald, mNeonGreen, or EGFP emission with MBS 488/561/639 and BP 420-480 + BP 570-630, BP 420-480 + BP 495-550, respectively. Z-stacks of 78.21 x 78.21 μm (1839 x 1839 pixels) were imaged for neurons expressing nNeonGreen-Synapsin and SiT-GFP. Imaging of Z-stacks of 66.32 x 66.32 μm (1560 x 1560 pixels) was performed for mEmerald-Sec61b. All Z-stacks were acquired using a piezo-motor and Nyquist-optimized z-step sizes (0.133 μm). Images were acquired and processed using automatic Airyscan processing in ZEN software (Zeiss).

#### Immunocytochemistry

For immunostaining experiments, cells were seeded in a lower density on poly-d-lysine–treated coverslips without cylinders. Cells were transfected with shRNA pLKO plasmid using Lipofectamine 2000 reagent or infected with shRNA pLKO plasmid carrying lentivirus. At DIV 16-18, cells on coverslips were fixed using 4% paraformaldehyde (PFA), treated with 50 mM NH4Cl, and then permeabilized for 10 minutes using 0.1% Saponin. Cells were blocked using a 5% bovine serum albumin buffer, and the antibodies were incubated in the same buffer. Anti-SLC13A5 antibody was used at 1:100 dilution, anti-TagBFP nanobody was used in 1:500 dilution, anti-Tuj1 antibody used in 1:1000 dilution. Alexa Fluor–conjugated fluorescent secondary antibodies were obtained from Life Technologies and used at 1:500. Cells were mounted on glass slides using ProLong Diamond (Thermo Fisher Scientific, P36970). For KD quantification in somas, neurons were sparsely transfected. SLC13A5 staining from regions of interested of the cell somas that were positive for nuclear BFP were compared to neighboring BFP negative cell somas. For axonal KD analysis, neurons were infected with control or SLC13A5 shRNA lentivirus, achieving almost 100% infection rate. Axonal KD efficiency was analyzed by selecting regions of interest that were Tuj1–positive as masks to compare SLC13A5 staining.

Immunocytochemistry for Airyscan microscopy imaging was done with following adjustments: Primary rat neurons were fixed in PBS (Fisher Scientific, 70-011-069) with 4% paraformaldehyde (Electron Microscopy Sciences, 50-980-487) and 4% sucrose (Fisher Scientific, S5-500) for 30 minutes at 37°C. Subsequently, cells were washed three times with PBS and incubated with 50 mM NH4Cl (Sigma Aldrich, 213462) in PBS for 10 minutes. After three washes with PBS (Fisher Scientific, 10010023), the cells were permeabilized with a 0.1% Triton-X100 solution (Sigma Aldrich, 93443) in PBS for 10 minutes. To prevent non-specific binding, a blocking step was performed for 30 minutes with a solution containing 5% BSA (Bovine Serum Albumin, Jackson ImmunoResearch Lab, Inc., 001-000-161) and 0.1% Triton-X100 in PBS. The primary antibody (anti-SLC13A5) was then added to the blocking solution (1:400), and the cells were incubated overnight at 4°C. After three rinses with 0.1% Triton in PBS for 10 minutes, the cells were incubated with the secondary antibody (Goat anti-Rabbit IgG (H+L) Highly Cross-Adsorbed Secondary Antibody, Alexa Fluor™ 568, ThermoFisher, A-11036) in blocking solution for 2 h. Finally, the cells were rinsed three times with 0.1% Triton in PBS for 5 minutes and three times with PBS for 5 minutes before being imaged in PBS. After imaging, the cells were then mounted in ProLong™ Diamond Antifade Mountant (ThermoFisher, P36961) for long-term storage.

### QUANTIFICATION AND STATISTICAL ANALYSIS

Images were analyzed using the ImageJ software. For time series analysis, ImageJ plugin Time Series Analyzer V3 was used. Statistical analysis was performed with GraphPad Prism v9. All statistical details can be found in figure legends. Unless stated otherwise, unpaired t-test was used for comparison of two groups, while ANOVA with Tukey post-hoc test was used to compare multiple groups. p < 0.05 was considered significant and denoted with a single asterisk, whereas p < 0.01, p < 0.001 and p < 0.0001 are denoted with two, three, and four asterisks, respectively. The n value, indicated in the figure legends for each experiment, represents the number of neurons imaged.

